# Daily turnover of active giant virus infection during algal blooms revealed by single-cell transcriptomics

**DOI:** 10.1101/2022.10.15.512338

**Authors:** Gur Hevroni, Flora Vincent, Chuan Ku, Uri Sheyn, Assaf Vardi

## Abstract

Viruses are the most abundant biological entity in the ocean and play a significant role in shaping the marine ecosystem. The past two decades have revealed an outstanding diversity of giant viruses infecting protists across the tree of life and, in particular, algae that form massive blooms in the ocean. Virus-induced bloom demise significantly impacts marine ecology and biogeochemistry, as well as the associated microbial community. Nevertheless, little is known about the infection dynamics of these giant viruses in the natural environment and their role in regulating algal blooms. Here, we provide evidence for a daily life cycle of giant viral infection in algal blooms by processing the transcriptome of over 12,000 single algal cells during different phases of interaction with their giant viruses. We revealed that viral infection occurs already at the exponential phase of the bloom and that the timing of infection can determine the magnitude of the bloom but not the fraction of infected cells. We further revealed that the same proportion of infected cells are in the early phase of the viral replication program (13.5%) throughout several consecutive days of the bloom, suggesting that a daily turnover of infection is at play during the bloom and demise phases of the algal population. This may imply that a continuous source of virocell-associated metabolites diffuses throughout the bloom succession and could fuel the microbial food webs. Finally, we link single cell infection state to host physiology and show that infected cells remained calcified even in the late stage of infection, contradicting common observation of bulk population in which viral infection is directly linked with decalcification. Together, these results highlight the importance of studying host-virus dynamics in natural populations at a single-cell resolution, which can provide a fresh view of the dynamics and propagation of viral infection. This approach will enable quantification of the impact of marine viruses on microbial food webs.

## Introduction

Algal blooms occur across oceans and lakes annually, covering up to thousands of square kilometers^1^. These hotspots of primary production impact global biogeochemical cycles^2,3^, release volatiles that contribute to cloud formation^4^, and influence marine microbial food webs^5^. Algal blooms can develop over weeks and are decimated within a few days due to a combination of environmental factors^6^. During bloom succession, phytoplankton cells can be subjected to abiotic stress (e.g. nutrient limitation) and biotic interactions with pathogens (e.g. viruses^7^, bacteria^8,9^, parasites^10^) or grazers^11,12^. A major limitation in evaluating mortality agents that can drive bloom dynamics is that most studies depend on bulk assessment of microbial abundances, thus averaging the entire population. This hinders any quantification of active pathogen infection on the subcellular level.

The coccolithophore microalga Emiliania huxleyi forms large oceanic blooms visible from space, and its calcite exoskeleton accounts for one third of the total marine CaCO3 production^13^. *E. huxleyi’s* specific giant double-stranded DNA virus (EhV) is regarded as the major mortality agent responsible for bloom termination^14–17^; its abundance can be assessed by quantification of extracellular virus-like particles (VLPs) via flow-cytometry^18^ or by qPCR using viral specific gene marker^19^. EhV identification can be complemented by transmission electron microscopy^20^ or other recent approaches based on tagging of viral DNA marker genes^21,22^. Bulk transcriptomics provide powerful insights on the host metabolic response and viral transcription program^23,24^, while coccolithophores optical properties teach us about host physiology such as *E. huxleyi’s* calcification and chlorophyll concetration^25^. However, we have not yet captured the importance of cell-to-cell heterogeneity in host response to viral infection that may influence bloom fate and ocean biogeochemistry.

Single-cell approaches offer a unique opportunity to zoom in into the infection dynamics and track an actively infected cell-the virocell^23,26^ - as opposed to only assess the outcome of infection, e.g. virion abundance in the extracellular milieu. Recent implementation of single-cell dual transcriptomics of *E. huxleyi* infection under controlled laboratory settings, revealed that viral infection follows a sequential transcriptional program^27^. Yet, basic questions remain unanswered: can we track single cell infection dynamics in natural blooms, and can we recapitulate the viral transcriptional program in nature? Resolving such processes in the natural environment will significantly improve our understanding of viral contributions to algal mortality and its possible implications on the flux and recycling of nutrients and metabolites^28^. This fundamental ecosystem process, termed the “viral shunt”, is a key driver of metabolic fluxes in the ocean and is responsible for fueling microbial life^7^. Thus, implementation of single cell approaches that enable assessment of the fraction of infected cells holds the promise to add a quantitative dimension to these ecosystem processes.

To characterize viral infection dynamics at a single cell resolution in natural populations, we conducted an induced bloom mesocosm experiment in Norway in May 2018. 11m^3^ of natural fjord water were enclosed in seven non-permeable bags, supplemented with nutrients, and monitored daily for 24 days^29^. A major bloom of *E. huxleyi* occurred in all bags (Figure S1), and we focused on three distinct bags that respectively showed low, medium, and high virus to host ratio, to conduct single cell transcriptomics analysis. We provide the first single cell view on the interplay between viral infection state and algal host physiology across thousands of cells at different phases of the bloom (Figure 1A). These results enabled quantification of the turnover of viral infection in individual cells of an algal bloom and highlight the importance of single-cell studies in addressing fundamental concepts of host-pathogen interactions in the marine ecosystem.

**Figure 1.**
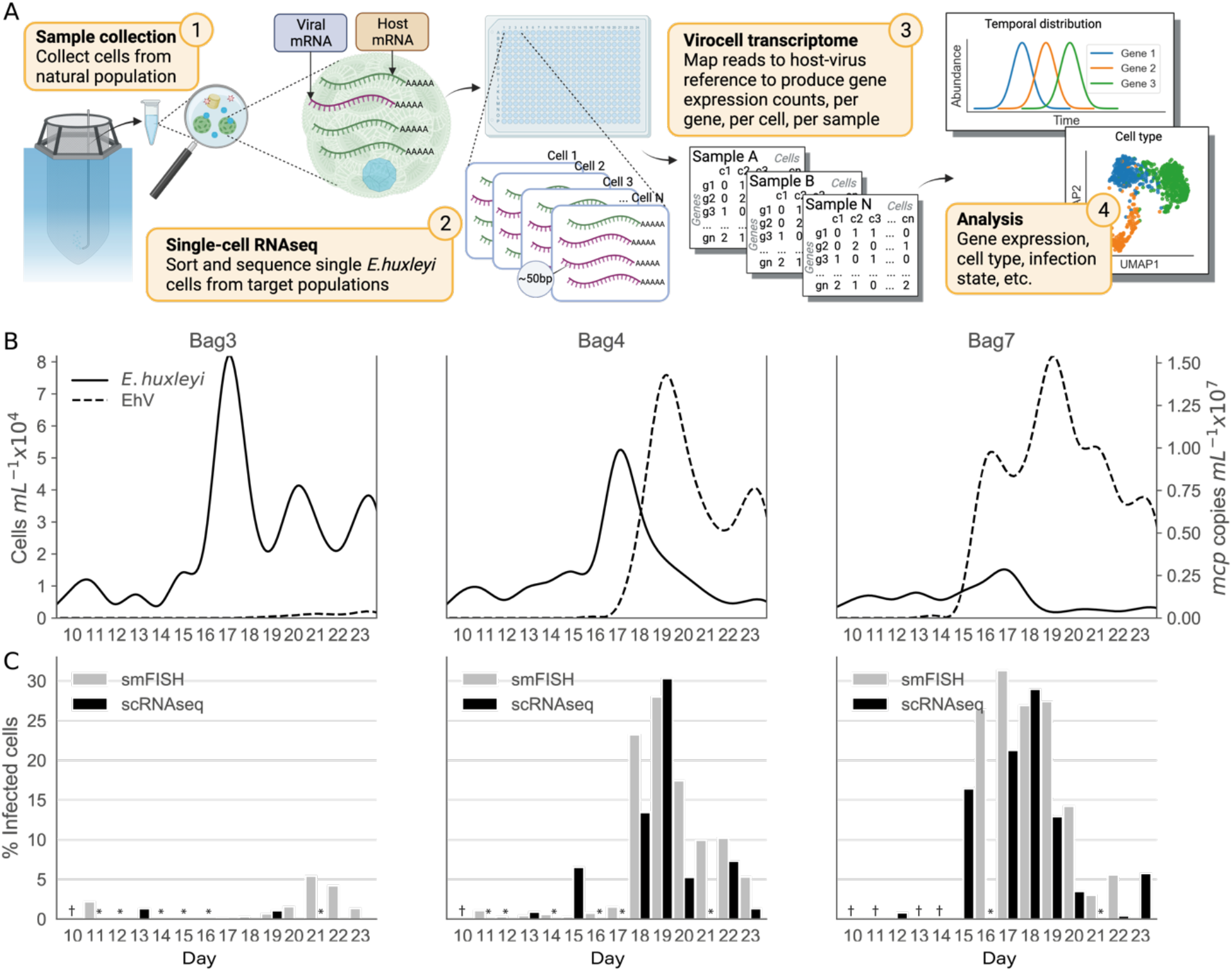
Dynamics of *E. huxleyi* and EhV infection in the mesocosm bags sampled for single cell analysis; (**A**) Schematic workflow of the single-cell sampling and analysis process: 1) sample natural populations from mesocosm bags. 2) sort single *E. huxleyi* cells and process for singe-cell RNAseq. 3) quantify single-cell expression based on dual host-virus transcriptome. 4) analyze host-virus interactions at the single-cell level; (**B**) *E. huxleyi* cell abundance based on flow cytometry analysis, and assessment of intracellular (2-20µm) viral abundance based on qPCR quantification of viral major capsid protein gene; (**C**) Quantification of percent of active viral infection based on single molecule FISH (smFISH) that assess the level of viral *mcp* transcript per cell (grey), and quantification of percent of infected cells based on total viral transcript (≥3 viral UMIs) per cell using single cell RNAseq (black); † represents time points for which no data was collected by either methods; * represents no data for one of the methods.

## Results and Discussion

### Early viral propagation determines bloom magnitude but not the fraction of infected cells

Throughout the bloom succession, *E. huxleyi* cell abundance, estimated by flow cytometry, increased in all bags around day 10, peaked at day 17, and rapidly declined from day 18 onwards (Figure 1B). Bag 3 reached the highest *E. huxleyi* abundance (8.1 x10^4^ cells/mL), followed by bag 4 (5.2 x10^4^ cells/mL) and bag 7 (1.5 x10^4^ cells/mL). The *E. huxleyi* virus (EhV) abundance, estimated by qPCR using the major capsid protein gene *mcp* on biomass associated filters (2-20 μm), reached a maximum of 1.54 x10^7^ *mcp* copies/mL and 1.42 x10^7^ *mcp* copies/mL in bag 7 and 4 respectively, and was mostly not detected in bag 3. At the peak of host abundance on day 17, this led to a higher virus to host ratio in bag 7 (∼500), followed by bag 4 (∼14) and finally bag 3 (∼0.02) (Figure S2). Substantial differences in viral infection dynamics were observed: in bag 4, EhV abundance peaked on day 19, two days after the host’s maximum abundance. In contrast, EhV abundance in bag 7 rapidly increased on day 16, before the host peak on day 17 (Figure 1B). As each mesocosm enclosure started with a similar inoculum, we hypothesize that these significant differences in viral dynamics are the product of small-scale stochastic effects, that can impact bloom succession into different states.

To gain higher resolution on viral infection, we used digital droplet PCR (ddPCR) for the *mcp* gene - more sensitive than qPCR^30^ – to quantify intracellular viruses using 2-20 μm biomass filters. We detected viruses as early as day 15, day 13 and day 9 in bag 3, bag 4, and bag 7 respectively (Figure S3). From the day of lowest difference between host abundance across bags (day 12, mean 5,306.3, sd 796.5 cells/mL), to the peak of the bloom (day 17), *E. huxleyi* abundance increased 18.6 folds in bag 3, 9.2 folds in bag 4, and only 2.5 folds in bag 7. Thus, the earlier infection was detected intracellularly, the lowest the host abundance fold change. In bag 4, the earliest occurrence of viral infection coincided with the mid exponential growth phase of the host (day 13). In contrast, viral infection in bag 7 was detected in early exponential growth phase of the host (day 9). This suggests that the timing of early infection, very early or during mid exponential growth phase, can decrease the maximal carrying capacity of the bloom, with possible impact on bloom succession and demise^29^. The detection of infection at very early stage of the bloom, namely at the beginning of the exponential growth phase in bag 7, suggests possible interplay between viral replication and growth of algal host. Indeed, although EhV is a lytic virus, it can also bud out of *E. huxleyi* cells during early phase of infection without leading to cell death^31–33^. This viral strategy can be of great advantage to increase encounter rates with growing host density at early stage of the bloom.

To assess the extent of viral infection within host cells, we used two independent methods to quantify intracellular viral transcription at the single-cell level, and calculated the percentage of actively infected cells in each day of sampling. The single-molecule FISH assay (smFISH) visualizes individual *mcp* transcripts in single cells and identifies *E. huxleyi* cells in natural microbial consortia using fluorescent probes targeting a specific 28S region^34^ (see methods). In bag 3, the fraction of infected cells was close to zero with a gradual rise to 5% towards the end of the experiment (days 21-22). In bag 4, it was previously shown that the proportion of infected cells increased dramatically within a day, from 1.5% on day 17 to 23.2% on day 18, peaking at 28% on day 19^33^. In bag 7, smFISH samples were available only from day 16 onwards, with a peak at day 17 of 31.3% of infected cells (Figure 1C).

Simultaneously, on several days during the bloom succession, more than 12,000 cells were sorted for dual single-cell RNAseq (scRNAseq) to follow the host and viral transcriptomes during active infection (Figure 1A). The sorted cells were gated based on their chlorophyll autofluorescence and side-scatter signal (a proxy for cell calcification level), separating the cells into three main groups: (1) Calcified *E. huxleyi* (high chlorophyll, high side-scatter) (2) Naked *E. huxleyi* (high chlorophyll, low side-scatter), and (3) Calcified low chlorophyll *E. huxleyi* (low chlorophyll, high side-scatter) (Figure S4). The sorted cells were processed using the MARS-Seq scRNAseq protocol^35^, and mapped to a dual reference containing both *E. huxleyi* and EhV genomes, as described in^27^. Briefly, cells were barcoded and each mRNA transcript received a unique molecular identifier (UMI). After sequencing, reads were mapped to the host and virus genome references, and UMI counts per gene and per cell were quantified (see Data Availability). This dataset constitutes, to our knowledge, the first dual host-virus single-cell transcriptome from environmental microbial samples. Previous lab data showed a small number of non-infected control *E. huxleyi* cells with random assignment of up to 2 viral UMIs^27^, hence, we only considered cells with 3 or more viral UMIs as infected cells. We quantified the percentage of infected cells and show that infected cells constituted a maximum of ∼30% from the total single cells sampled per day (Figure 1C), thus supporting the smFISH results. We cannot rule out the possibility of a higher proportion of infected cells at a time that was not sampled. However, this is unlikely based on the relatively high frequency at which the samples were collected and the similar pattern between multiple quantification methods (qPCR, smFISH, scRNAseq). The two independent measures of active infection at the single-cell level (smFISH and scRNAseq) indicate that viral infection contributes up to a third to host mortality in the infected bags, suggesting that other mortality agents can lead to *E. huxleyi* bloom demise, such as protists grazing^11,12^, pathogenic bacteria^8,9^, or programmed cell death induced by signaling molecules released by infected cells^36^.

### Studying the life cycle of viral infection at a single cell resolution in algal blooms

One major question arising from intracellular viral infection dynamics, is whether the maximum of 30% of infected cells detected on days 17-19 represents an accumulation of cells from previous days or is formed by a daily turnover of infection preventing 100% of infected cells at steady state. To this end, we clustered the single cells based on their viral transcriptional profile (Figure 2A), to determine their position along a viral infection continuum. Viral genes were previously annotated according to their sequential transcription during infection in lab cultures, and grouped in kinetic classes (*Immediate-early, Early, Early-late, Late1, and Late2*) to define each viral infection state^27^. Visualizing the total gene expression by kinetic class within natural samples revealed a distinct population of single-cells (Cluster 0) expressing genes in the early kinetic classes like *Immediate-early*, and *Early* (Figure 2B-F). 63% of the transcripts in Cluster 0 came from *Immediate-early* genes, 25% from Early genes, and only 8% from Late2 genes. In cluster 1, 18% of the transcripts came from Immediate-early genes, 51% from Early genes, 15% from *Early-late* gene, and 16% from *Late2* genes. In contrast, cells in clusters 2-5 are dominated by transcripts from *Late2* genes (68-81% of the cluster) and have similar proportions of kinetic class distribution (Figure 2G). The abundance of *Immediate-early* transcripts thus drastically decreased between cluster 0-1 and cluster 2-5 (Figure 2G) similar to laboratory experiments of a time course of infection of *E. huxleyi* cells (Figure 2H, data from^27^).

**Figure 2.**
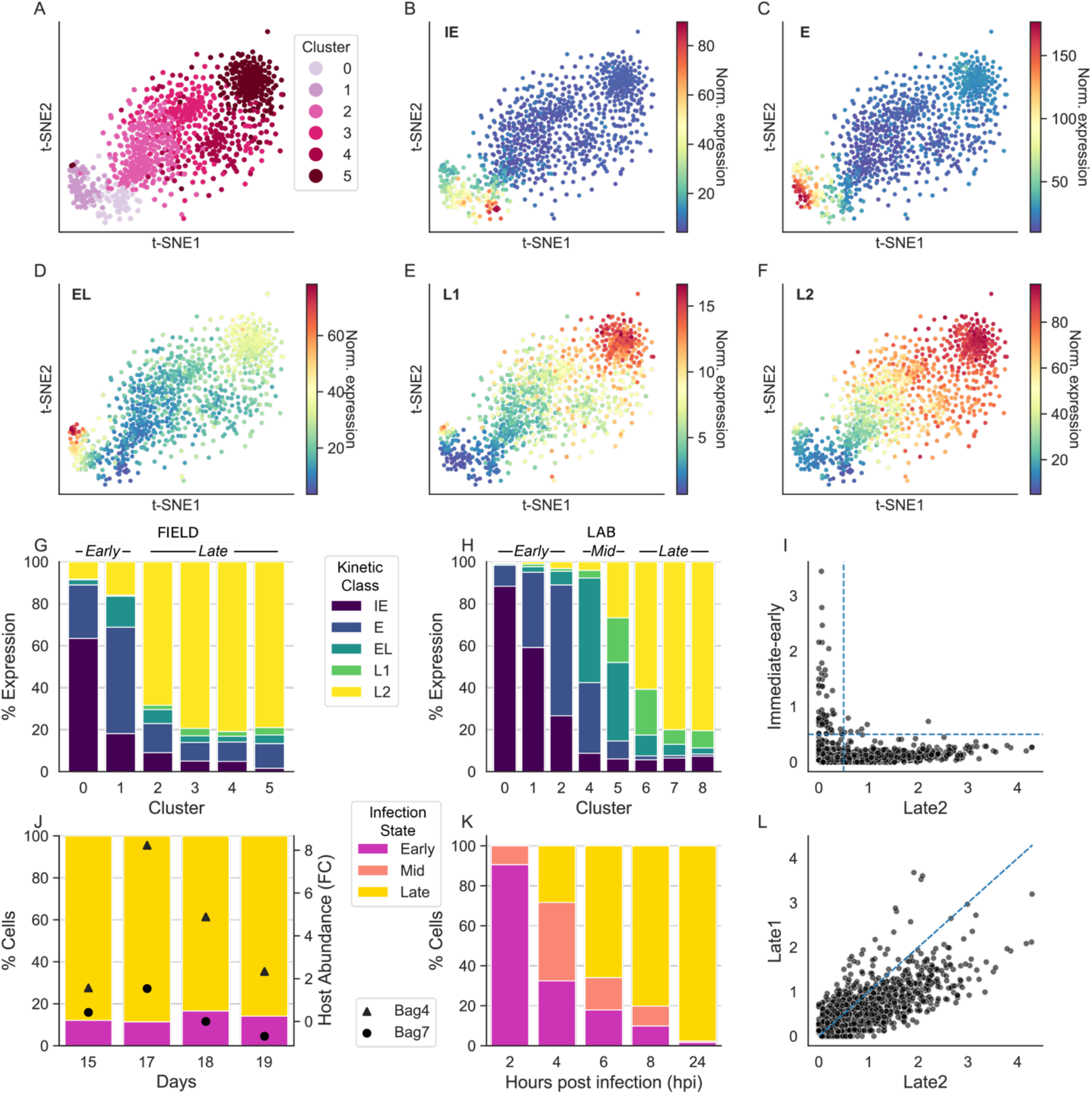
Daily turnover of the infection cycle observed by analysis of viral kinetic class expressed in single cells during a bloom. (**A**) t-SNE plot of infected cells (n=1,038; minimum 10 viral UMIs per cell) collected from bags 4 and 7. Each dot is a single cell and the color represents the cell’s assigned cluster. (**B-F**) t-SNE plot of gene expression summed by viral kinetic class: (**B**) *Immediate-early*; (**C**) Early; (**D**) Early-late; (**E**) Late1; (**F**) Late2; (**G**) Percentage of kinetic class expression by cluster for mesocosm cells; and (**H**) lab cells; (**I**) Scatter plot of the expression of *Immediate-early* vs. Late2 genes, and (**L**) Late1 vs. Late2 genes in the infected cells; (**J**) Percentage of cells by infection state, Early, Mid, Late by day for mesocosm cells; and (**K**) by hour for lab cells. The triangular and circular markers represent the fold-change in host abundance during bloom and demise in the mesocosm, compared with Day12 (**J**).

To better resolve the progression of viral infection, we defined cells in clusters 0 and 1 in the natural population samples as cells in Early infection state, and cells in clusters 2-5 as cells in Late infection state (Figure 2G). This separation is supported by the expressions of *Immediate-early* and Late2 genes per cell (the two extremes in the viral infection program) that show a mutually exclusive expression pattern (Figure 2I), while Late1 and Late2 are mostly co-expressed (Figure 2L). This pattern implies that the transcriptional viral program is sequential in natural populations similar to that observed in the lab. Under this pattern, early viral genes are downregulated as the viral program transitions to the late infection phase. Interestingly, levels of *Immediate-early* and Late2 gene expression could be used as marker genes to rapidly determine a cell’s infection state in oceanic blooms using bulk metatranscriptomic analysis (see Table S1 for a list of potential viral infection state markers). As a result of the progressive gene expression during viral infection, we expected to find higher proportion of cells in Early state at the beginning of the bloom, and enrichment of cells in Late state during the demise. Surprisingly, when plotting the proportion of cells by infection state by day, covering the transition from exponentially growing population (days 15-17) to its decline (days 18-19), we found almost no difference in the proportions of Early and Late cells between days (average 13.5%, sd 2.3% for cells in Early infection state. Figure 2J). The sequential nature of the EhV infection program and the constant daily proportions of infection states suggests that a daily turnover of the infection cycle is at play in natural populations. Another scenario to explain the constant ratio is if cells remain in the same kinetic class for several consecutive days. However, based on lab data we find such an option highly unlikely as a very rapid transition occurs from early to late infection. Using the same clustering approach on the dataset generated during viral infection of *E. huxleyi* cultures in the lab, we defined cells in clusters 0-2 as Early, cells in clusters 4 and 5 as Mid infection state (Mid), and cells in clusters 6-8 as Late (Figure 2H). Lab cultures of simultaneously infected (high MOI) single cells showed 91% of the population was composed of Early cells and 9% of Mid cells 2 hours post infection (hpi), and as early as 4 hpi infection 68% of cells transitioned to Early state. By 8 hpi 90% of the cells were in either Mid or Late infection states (10% and 80% respectively. Figure 2K), which illustrates the very short residential time of cells in early infection state.

The notion of a daily EhV infection cycle in the natural environment is supported by other studies in photosynthetic microorganism, where viral infection cycles have shown temporal association with diel cycles and photosynthesis-related host metabolism in bulk measurements^37–39^. This has important implications for our understanding of bloom dynamics. First, it shows that the exponential growth phase of a bloom, classically described as a “healthy” physiological state on the population level, is composed of a subpopulation of cells that are already in late phase of viral infection. This decoupling between the fate of the individual cell as compared to bloom fate can only be now unraveled by using single cell analysis. Second, the proportion between these co-existing phenotypes e.g., susceptible cells, virocells at different infection states, and resistant cells, will eventually determine the ecological and biogeochemical outcome of the bloom. Each phenotype displays a different metabolism, such as *E. huxleyi*/EhV virocells who produce unique viral glycosphingolipids^36^, that can differentially affect the neighboring microbiome upon cell lysis. A daily turnover of viral infection thus implies that a quotidian viral shunt is at play, whose absolute impact is driven by host cell abundance.

### The interplay between host physiology and infection states

Given the phenotypic heterogeneity in the extent of viral infection among individual cells, we sought to examine the interplay between infection states and host physiology within *E. huxleyi* cells. High calcification and chlorophyll levels are often considered as healthy physiological states, serving as proxies for *E. huxleyi* intact calcium carbonate exo-skeleton, and active photosynthesis. Viral infection has many consequences on host physiology, in particular decalcification^25^, for which the reasons are still subject to debate^40^. To test this, we used indexed sorting FACS data which records each cell’s forward scatter (a proxy for size), side scatter (a proxy for calcification), and chlorophyll autofluorescence values before their sorting and downstream single-cell transcriptomics analysis.

For the following analysis we used data collected from bag 4 in day 19 in which the level of infection was approximately 30% in both smFISH and scRNAseq assays (Figure 1C). The non-infected cells are almost evenly distributed between the chlorophyll vs. side-scatter gates (Figure 3A). However, most of the infected cells, whether in Early (72.94%) or Late (79.82%) stages of infection, fell within the high calcified *E. huxleyi* flow cytometry gate (high-chlorophyll, high-side-scatter) (Figure 3B-C). This observation stands in contrast to the general notion of a direct link between viral infection and the shedding of coccoliths by the host cell, a scenario in which most of the cells in Late stage should thus appear in the lowly calcified gate. One hypothesis is that viral infection enhances continuous coccolith production and shedding, as was recently suggested based on independent measurements of enhanced production of particulate inorganic carbon^29^. The phase of continuous shedding can be followed by decalcification in the bloom decline, associated with general cell death processes occurring at the lytic phase, after viral egress from the cells^31,33^. This observation also highlights the importance of single cell approaches in assessing the consequences of viral infection on host physiology.

**Figure 3.**
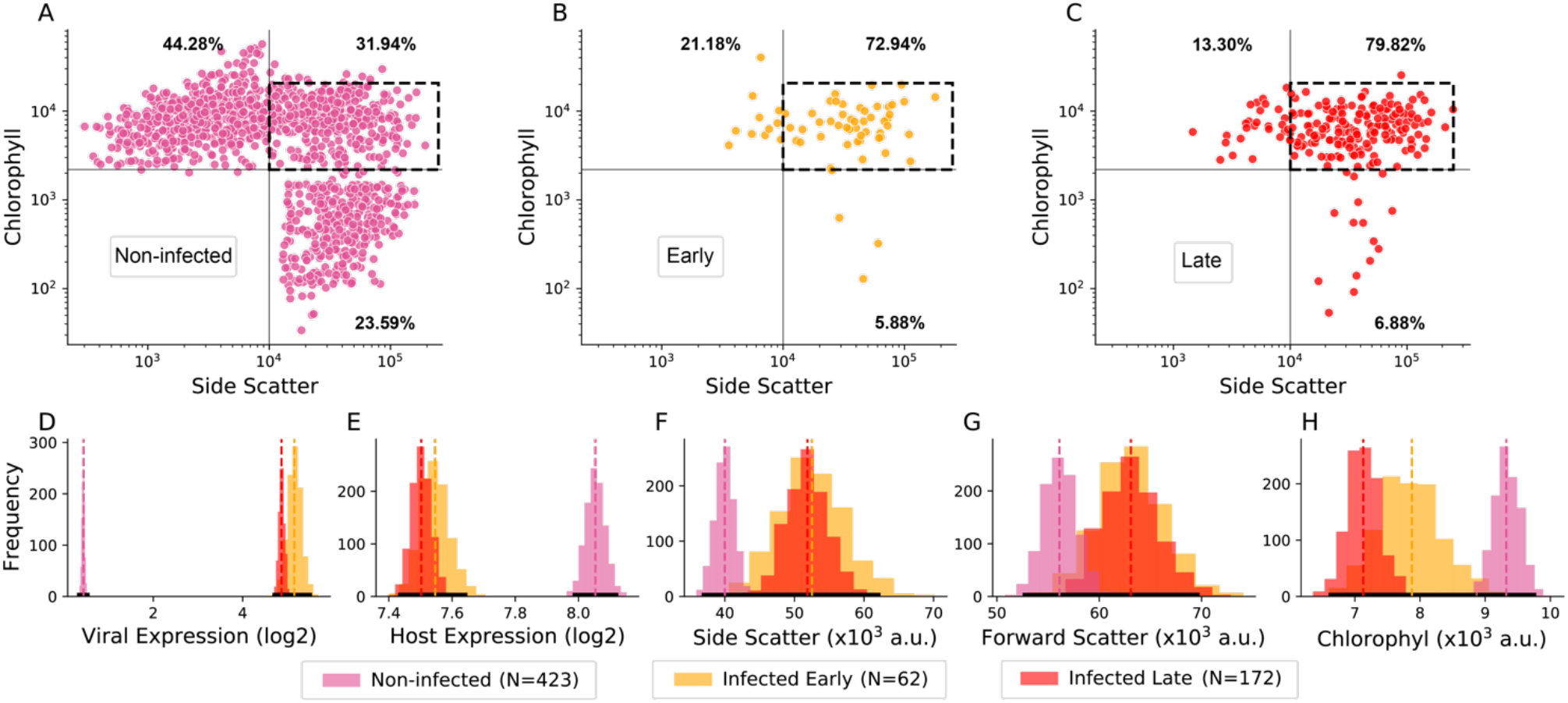
The interplay between physiological and transcriptional states at a single cell resolution during viral infection. (**A-C**) Scatter plot of sorted cells by infection state. Cells are positioned according to their side scatter (x axis) and chlorophyll (y axis) values recorded by index sorting flow-cytometry. (**A**) non-infected cells; (**B**) Early infection state; and (**C**) Late infection state; (**D-H**) Distributions of the mean values (bootstrapped) of intra- and extracellular properties for the three subpopulations (Early, Late and non-infected cells) from the calcified *E. huxleyi* sorting gate (dashed quadrant in panels A-C). (**D**) Viral expression; (**E**) Host expression (**F**) Side scatter (**G**) Forward scatter and (**H**) chlorophyll.

To further identify differences between the infected and non-infected subpopulations, we focused on cells from the calcified *E. huxleyi* gate (Figure 3A-C dashed rectangular). To this end, we looked at the 95% confidence interval (CI) around the mean values distribution of both molecular (gene expression) and physiological (indexed sorting) parameters within the three groups of non-infected, Early and Late infected cells (see methods). As expected, the viral gene expression was significantly higher in the infected populations compared to non-infected cells (Early C.I. [4.86, 5.47], Late C.I. [4.75, 5.00], non-infected C.I. [0.37, 0.49]. Figure 3D). The host expression was significantly lower in infected cells (Early C.I. [7.45, 7.64], Late C.I. [7.44, 7.57], non-infected C.I. [7.99, 8.11]. Figure 3E) suggesting that the cellular transcription is undergoing a shutoff upon infection. The side scatter values, a proxy for the level of calcification and cellular granularity, were significantly higher in the infected population (Early C.I. [43.09, 61.86], Late C.I. [45.89, 57.80], non-infected C.I. [37.19, 42.74], Figure 3F). There was no significant difference in the forward-scatter between infected and non-infected cells (Figure 3G). The chlorophyll values were significantly lower in the infected cells (Early C.I. [6.89, 8.80], Late C.I. [6.65, 7.62], non-infected C.I. [8.94, 9.73]. Figure 3H). The co-existence of infected and non-infected cells within the gate of calcified *E. huxleyi* emphasizes the knowledge gap stemming from studying host-virus dynamics on the bulk-level, where cell-cell heterogeneity is averaged out.

To further investigate the host response, we pooled all single-cell data from days 15, 17, 18, 19 and 20; most infected cells were detected in those samples that cover the bloom and demise phases. We selected cells with at least 100 host UMIs or 10 viral UMIs (3,996 cells), clustered them based on their expression profiles including both host and viral genes (Figure 4A) and show distinct clustering of non-infected (0-3) and infected (4-5) cells. Cluster 4 was dominated by cells expressing viral genes of early kinetic classes (*Immediate-early*, Early, and Early-Late; 156 cells. Figure 4B); while in cluster 5 Late viral genes dominated (Late1 and Late2; 686 cells. Figure 4C). Cluster 0-3 represented a majority of cells that are yet be infected but still can be primed by exposure to infochemicals and vesicles produced by infected cells. Cells in cluster 0 generally showed higher expression levels of genes related to glycolysis (Figure 4D), cells in cluster 2 had higher expression of genes related to nucleotide synthesis (Figure 4E) and cells in cluster 3 were enriched in genes related to sphingolipids biosynthesis (Figure 4F). Differential expression of genes within those clusters could be a consequence of position in the cell cycle that we were not able to resolve. Overall, host transcription was mostly downregulated in infected cells with a rapid transition from host to viral transcriptome (Figure 4G, H); as part of the viral takeover strategy. The alignment of the rapid cellular shutoff corroborates observations from viral infection dynamics in cultures, suggesting that the rapid viral hijacking strategy is also predominant in viral infection in natural *E. huxleyi* blooms.

**Figure 4.**
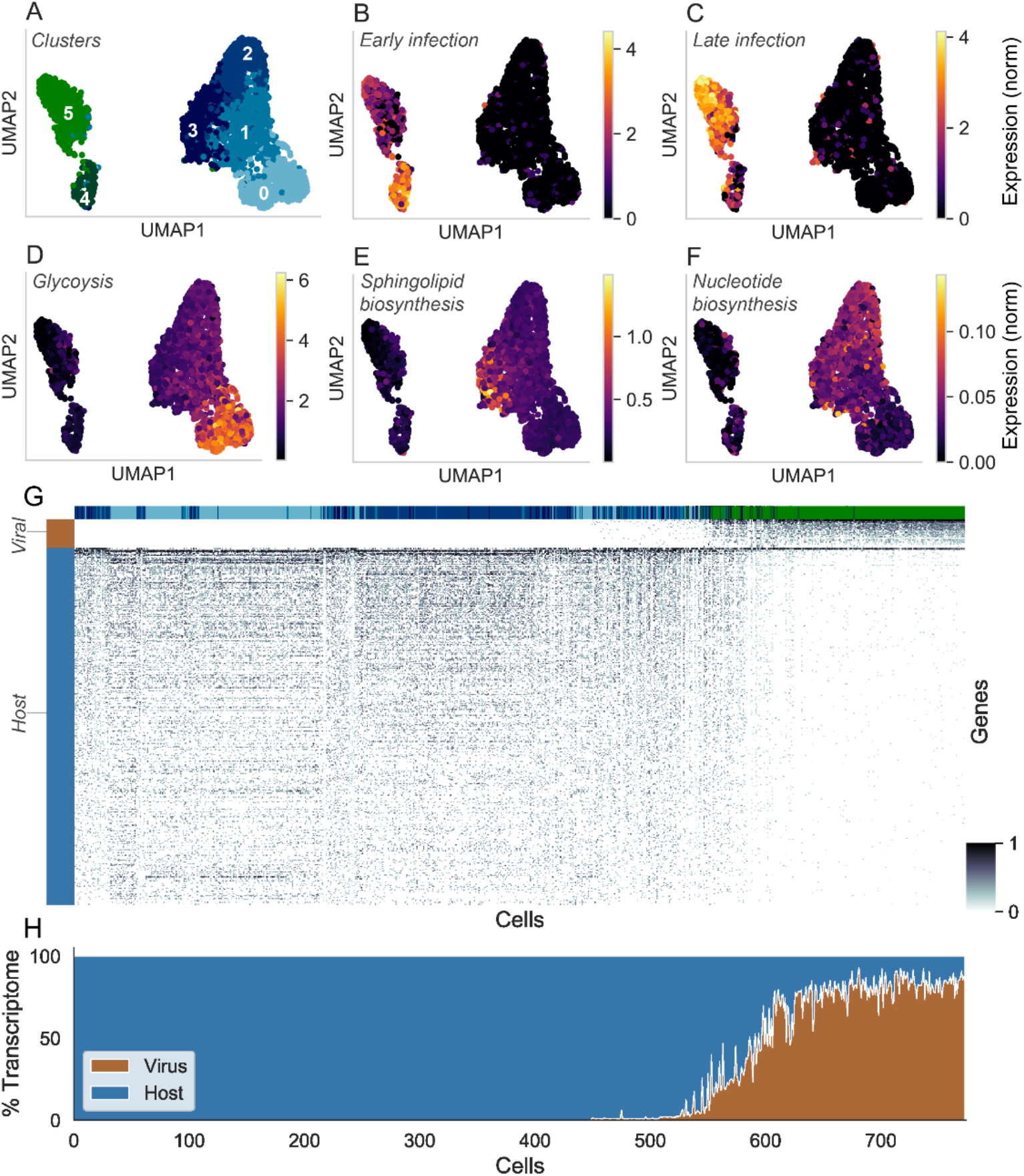
Host response to viral infection resolve on a single cell level. (**A**) Uniform Manifold Approximation and Projection (UMAP) plot of single cells clustered according to host-virus dual transcriptome (N=3,996). Each dot is a cell and the color represent the cell’s cluster affiliation; (**B**) Viral early and (**C**) late kinetic class gene expression (log) overlayed on the single-cell UMAP plot; Host expression of pathway related genes for (**D**) Glycolysis; (**E**) Sphingolipid biosynthesis; (**F**) Nucleotide biosynthesis; (**G**) Heat map of genes (rows; color bar: host-blue, virus-brown) expressed in each cell (columns; color bar: clusters from A); (**H**) Percentage of expression of viral and host genes in the single-cell transcriptomes.

The very rapid host shutoff constitutes a challenge in the quest of better identifying host genes that could potentially be upregulated during infection at the single cell level. These genes can be involved in defense pathways that are still yet to be discovered in marine algae, given the observation of a rare *E. huxleyi* subpopulation (<1%) that can recover from infection^41^. We employed different methods to detect induction of host genes in relation to viral transcription, including pairwise gene-gene relationship and differential expression analyses. Unfortunately, the low level of host gene expression in infected cells limited the significance of the results, that could be improved by better recovery of single-cell transcriptomes (for example CELL-Seq^42^) and sampling of more cells or dedicated experimental design. It can also help to unravel the upregulation of cellular processes that potentially benefit viral replication such as nucleotide synthesis to meet the high demands of large giant virus genomes, glycosphingolipid synthesis that are necessary for virions assembly, or general stress response including production of reactive oxygen species^27,43,44^. Several challenges remain to be solved given that our data originates from complex natural populations, such as the variability of host and viral strains, a lower frequency of sampling (daily vs. hourly in the lab), and the impact of multiple uncontrolled variables like temperature, nutrients level, and other interacting species.

## Conclusion

By applying single cell dual host-virus transcriptomics to natural populations, we were able to quantify with unprecedented resolution active viral infection within an algal bloom. Surprisingly, we observed a daily turnover in viral progression at the single cell level, despite synchronized algal bloom and demise of the population. This suggests that, throughout the bloom, there are cells that continuously become infected and lyse. Based on this evidence of persistent viral-derived cell lysis, we propose an alternative view on the consequences of viral infection in the marine ecosystem, in which a quotidian viral shunt is at play during algal blooms. Daily infection cycles of algal cells can result in a constant release of metabolites characteristic of the virocell metabolism, that can trigger a cascade of ecological and biogeochemical events. These include continuous fueling of a specialized eukaryotic and bacterial microbiomes^7,29,45^, induction of cell death in neighboring non infected cells^36^, improved growth rate of predators^46^, and even daily emissions of volatiles into the atmosphere^47^. Overall, this fine-grained assessment of host-virus interactions in marine ecosystems sheds new light on viral infection dynamics and how it can drive the viral shunt in the marine ecosystem.

## Data Availability

The documentation of the data used in this paper, including the raw UMI tables generated from the single cell MARS-Seq analysis are available in Dryad (https://doi.org/10.5061/dryad.g4f4qrftf). The Python code for processing data and generating figures is available at https://github.com/vardilab/daily-turnover-of-giant-virus-infection

## Materials and Methods

### Mesocosm core setup

The mesocosm experiment AQUACOSM VIMS-Ehux was carried out for 24 days between 24th May (day 0) and 16th June (day 23) 2018 in Raunefjorden at the University of Bergen’s Marine Biological Station Espegrend, Norway (60°16′11N; 5°13′07E). The experiment consisted of seven enclosure bags made of transparent polyethylene (11 m^3^, 4 m deep and 2 m wide, permeable to 90% photosynthetically active radiation) mounted on floating frames and moored to a raft in the middle of the fjord. The bags were filled with surrounding fjord water (day −1; pumped from 5 m depth) and continuously mixed by aeration (from day 0 onwards). Each bag was supplemented with nutrients at a nitrogen to phosphorus ratio of 16:1 according to the optimal Redfield Ratio (1.6 μM NaNO_3_ and 0.1 μM KH_2_PO_4_ final concentration) on days 0-5 and 14-17, whereas on days 6, 7 and 13 only nitrogen was added to limit the growth of pico-eukaryotes and favor the growth of *E. huxleyi* that is more resistant to phosphate limited conditions.

### Flow Cytometry measurements

Samples for flow cytometric counts were collected twice a day, in the morning (7:00 a.m.) and evening (8:00-9:00 p.m.) from each bag and the surrounding fjord, which served as an environmental reference. Water samples were collected in 50 mL centrifugal tubes from 1 m depth, pre-filtered using 40 μm cell strainers, and immediately analyzed with an Eclipse iCyt (Sony Biotechology, Champaign, IL, USA) flow cytometer. A total volume of 300 μL with a flow rate of 150 μL/min was analyzed. A threshold was applied based on the forward scatter signal to reduce the background noise.

Phytoplankton populations were identified by plotting the autofluorescence of chlorophyll versus phycoerythrin and side scatter: calcified *E. huxleyi* (high side scatter and high chlorophyll), Synechococcus (high phycoerythrin and low chlorophyll), nano- and picophytoplankton (high and low chlorophyll, respectively). Chlorophyll fluorescence was detected by FL4 (excitation (ex): 488nm and emission (em): 663-737 nm). Phycoerythrin was detected by FL3 (ex: 488 nm and em: 570-620 nm). Raw.fcs files were extracted and analyzed in R using ‘flowCore’ and ‘ggcyto’ packages and all data is available on Dryad^48^.

### Enumeration of extracellular EhV abundance by qPCR

DNA extracts from 2-20 μm filters from the core sampling (described extensively in^29^) were diluted 100 times, and 1 μL was then used for qPCR analysis. EhV abundance was determined by qPCR for the major capsid protein (*mcp*) gene: 5′-acgcaccctcaatgtatggaagg-3′ (mcp1F) and 5′-rtscrgccaactcagcagtcgt-3′ (mcp94Rv). All reactions were carried out in technical triplicates using water as a negative control. For all reactions, Platinum SYBER Green qPCR SuperMix-UDG with ROX (Invitrogen, Carlsbad, CA, USA) was used as described by the manufacturer. Reactions were performed on a QuantStudio 5 Real-Time PCR System equipped with the QuantStudio Design and Analysis Software version 1.5.1 (Applied Biosystems, Foster City, CA, USA) as follows: 50°C for 2 min, 95°C for 5 min, 40 cycles of 95°C for 15 s, and 60° C for 30 s. Results were calibrated against serial dilutions of EhV201 DNA at known concentrations, enabling exact enumeration of viruses. Samples showing multiple peaks in melting curve analysis or peaks that were not corresponding to the standard curves were omitted. Data is available in^48^.

### Enumeration of extracellular EhV abundance by ddPCR

Digital droplet PCR (Bio-Rad, Hercules, USA) was performed on 0.2 and 2 μm mesocosm filters of each bag including the fjord, to assess the absolute concentration of EhV. EhV abundance was determined using primers targeting the major capsid protein (*mcp*) gene: 5′-acgcaccctcaatgtatggaagg-3′ (mcp1F) and 5′-rtscrgccaactcagcagtcgt-3′ (mcp94Rv). Sample mix consisted of 10 μL of 2X QX200 ddPCR EvaGreen supermix, 1 μL of 2uM forward primer, 1 μL of 2 μM reverse primer, 5 μL of water and 5 μL of the DNA sample. To load the optimal amount of DNA, DNA extractions were diluted 1:10 and DNA concentration was measured using a Qubit dsDNA HS Assay Kit (Invitrogen, Waltham, USA). Depending on the concentration, between 1-5 μL of extracts were completed to a total of 5 μL with ultra-pure water and used in the final ddPCR reaction. Less than 80 ng of DNA was used for each reaction. From the final mix of 22 μL, 20 μL of each sample were loaded in the DG8 Cartridge and inserted in the QX200 droplet generator. Each cartridge contained a negative control containing the ddPCR mix with 5 μL of water. After droplet generation, samples were transferred to a 96 well-plate and inserted in a C1000 Touch thermal cycler. The following cycle was used: 95°C 5 min, followed by 40 cycles of 96°C for 30 sec, 58°C for 1 min, 4°C 5 min, 90°C 5 min and infinite hold at 4°C. After thermal cycling, the 96-well plate was read in the QX200 Droplet Reader and results analyzed using the Quantasoft software.

Quantasoft provides a final concentration of target copies/μL of ddPCR reaction. For mesocosm samples, we first calculated the total amount of target copies in 20 μL of ddPCR reaction and normalized it by the amount of sea water that was sampled, to obtain a final concentration of target copies/mL of sampled sea water.

### Quantification of infected cells using smFISH in natural samples

Sample fixation, storage and initial chlorophyll washes of field samples was identical to laboratory samples as described in^33^. In order to identify *E. huxleyi* cells, we designed a single probe specific to the 28S region of *E. huxleyi*^34^ (EG28-03, 5′-TAAAGCCCCGCTCCCGGGTT-3′, bound to C3-Fluorescein (ex/em = 490/525nm). Two helper probes were used (Helper A, 5’-GCCAGGACGGGAGCTGGCCG-3’ and Helper B, 5’-GAGGCGCGGCGCCGAGGCGC-3’). 28S, HelperA, HelperB were used at final concentration of 0.5μM, at the same time as the *mcp* probes at 0.1ng/mL final concentration, that target the *mcp* mRNA as a proxy for active intracellular infection. The helper probes facilitate unfolding of the 28S rRNA molecule by binding to the flanking regions of the probe of interest EG28. All samples were stained in 50μL of 40% formamide hybridization buffer, incubated at 37 degrees overnight. Samples were acquired using the ImageStreamX MarkII machine (ISX, Amnis, Luminex). Four excitation wavelengths were used: 405 nm for the DAPI (DAPI - Channel 7-50 mW), 488 nm for 28S rRNA (AF488 - Channel 2 - 200mW), 561 nm for *mcp* (TMR - Channel 3 – 200 mW). Data was analyzed using IDEAS6.2 (Amnis, Luminex). The compensation matrix was built using the IDEAS wizard and manually checked, before being applied to all the acquired files. Based on the Area (the number of microns squared in a mask) and Circularity (the degree of the mask’s deviation from a circle) of DAPI three populations are identified as single cells (mainly DAPI area < 60 a.u), doublets and aggregates (mainly DAPI area > 60 a.u). Single cells were additionally selected in the same focal plane using the Bright Field gradient and contrast (both gradient and contrast measure the sharpness quality of an image by detecting large changes of pixel values in the image). All gates were defined on a single file before being applied the total data set. Each file was then manually inspected to check the accuracy of single cell and aggregates gating. All the data (fluorescent intensities, morphological features, populations) was then exported for each cell of each file for analysis in R.

### Single-cell isolation by FACS and massively parallel scRNA-seq

The scRNA-seq method is based on the MARS-seq2.0 protocol^35^. Prior to the mesocosm experiment, 384-well plates were prepared using a Bravo liquid handling platform (Agilent) to transfer into each well a 2-μl lysis buffer containing 2 nM reverse transcription (RT) primers with cellular and molecular (UMI) barcodes. Plates with cell barcoding and lysis buffer were prepared at the Weizmann Institute and sent on dry ice to the Flow Cytometry Core Facility of the Haukeland University Hospital of the University of Bergen prior to the mesocosm experiment. During the experiment, 200mL of water were collected from individual mesocosm bags and prefiltered on a 20 μm mesh, kept in the dark in a cool box and sorted on the same day at the Haukeland Hospital. Single cells were isolated into individual wells in 384-well plates on a FACSAria II (BD Biosciences) using a 488-nm laser for excitation and a nozzle size of 100 μm. Cells were identified based on chlorophyll red autofluorescence (emission 663 to 737 nm) and separated under the “Single Cell” mode for maximal purity. Plates with sorted cells were immediately frozen on dry ice and later stored at −80°C before being sent back to the Weizmann Institute.

After thawing, the plates were heated to 95°C for 3 min to evaporate the lysis buffer, cooled, added RT reagent mix, and placed in a Labcycler (SensoQuest) for RT reaction. The unused primers were then degraded with Exonuclease I (NEB). cDNA from each set of 384 wells (one pool per plate) underwent second-strand synthesis and in vitro transcription to amplify the sequences linearly. The RNA products were fragmented and ligated with single-stranded DNA adapters containing plate-specific barcodes. A second round of RT was carried out, followed by polymerase chain reaction with primers containing Illumina adapters to construct DNA libraries for paired-end sequencing on a NextSeq machine (Illumina).

### Read processing and mapping to reference sequences

The FASTQ files were processed using the analytical pipeline of MARS-seq2.0^35^, which mapped the reads to viral and host reference sequences and demultiplexed them based on the pool, cellular, and molecular barcodes. At least four reads per UMI were obtained per cell. For the virus, the predicted CDSs in the EhV201, EhV163, EhVice and EhVM1 genome sequence^49^ were used as reference. For the host, an integrated transcriptome reference of *E. huxleyi*^50^. In addition to the nuclear sequences, the transcriptome reference contains chloroplast and mitochondrial transcripts, which were identified by the basic local alignment search tool (BLAST)^51^ searches against the respective organellar genome sequences^52^.

### Preprocessing of viral and host transcript abundance

Cells with zero UMIs (including all viral and host transcripts) as well as cells with the lowest 5% number of UMIs (i.e., <5%) were removed for most downstream analyses. Cells with the highest 5% number of UMIs (i.e., >95%) were also removed to prevent cases of wells with doublet or multiplet cells. The raw UMIs were further preprocessed using the Python package scprep version 1.0.10: Low expressing genes were filtered with *filter.filter_rare_genes and min_cells=5*, expression was normalized by cell library size with *normalize.library_size_normalize*, and the data was scaled with *transform.sqrt*.

### Infected cells clustering and visualization

Cells with 10 or more viral UMIs (infected cells) from mesocosm bag 4 and bag 7 were grouped into a single dataset with 1,038 cells and 220 genes. To visualize the cells in a two-dimensions based on their gene expression profiles, t-distributed stochastic neighbor embedding (t-SNE) dimensionality reduction was performed using the *TSNE* method in the *manifold* package of the Python library scikit-learn version 0.24.1 with perplexity=30. The infected cells were clustered using a Python package of the unsupervised clustering algorithm phonograph^53^ with default parameters.

### Viral kinetic class quantification and infection state definition

Viral gene UMIs were grouped and aggregated by their kinetic class assignment as defined in Ku et al.^27^ (Immediate early, Early, Early-Late, Late1, and Late2). The aggregated kinetic class gene expression was visualized per cell and per cluster. The infection state of cells was defined cluster affiliation. Cells in clusters with more than 50% expression of *Immediate-early* and Early kinetic class were defined in *Early* infection state. Cells on clusters with more than 50% expression of Late1 and Late2 kinetic class were defined as *Late* infection state. And cells in clusters with neither of the above states (either category below 50%) were defined as *Mid* infection state. The cells were then grouped by their infection state and visualized over time.

### Visualizing and bootstrapping single-cell physiological proxies

The indexed sorting FACS data was records provide physiological proxies for cells size (forward scatter), level of calcification (side scatter), and chlorophyll content (chlorophyll autofluorescence) per cell. These proxies were visualized with a scatter plot (side-scatter vs. chlorophyll) to detect cells populations by their physiological state, in conjunction with their infection state as defined by the single-cell transcriptomic data. To detect statistical differences between infection states of populations with seemingly similar physiological state (i.e., calcified *E. huxleyi*), N cells were randomly sampled 1000 times with replacement from each infection state group (where *N=group size*), and for each sample the mean was calculated for the different physiological proxies, per group (Early, Late, and non-infected). A significant difference between the distribution of the means of the different groups was determined by non-overlapping 95% confidence intervals (CI), calculated using the stats package of the Python library scipy version 1.5.3.

### Dual host-virus transcriptome analysis

A total of 3,996 cells with at least 100 host UMIs or 10 viral UMIs were selected for the dual transcriptome analysis of intracellular interactions between the host and the virus transcription. The raw UMI counts of the selected cells were further processed with the Seurat R package ^54^ version 4.0.0. The UMI counts were transformed using the SCTransform, scaled using the ScaleData with default parameters. The cells dimensionality was reduced using RunPCA (*npcs=50*), and with *RunUMAP* (*reduction=“pca”, dims=1:50*). Cells were clustered using FindNeighbors (*reduction=“pca”, dims=1:50*), and FindClusters (*resolution=range(start=0.4,end=0.9,step=0.1*)). The final clustering resolution was set to 0.4 based on visual inspection, resulting in 6 clusters (0-5). Differentially expressed markers between clusters was detected using *FindAllMarkers* with the MAST test^55^. For further analysis and visualization, the cells gene expression was aggregated based on host genes (by total expression, or by selected metabolic pathway marker genes), and viral genes (by total expression, or by kinetic class marker genes).

## Aknowledgements

We thank Brith Olsen Theil Bergum from the University of Bergen for her help and assistance in the Flow Cytometry Core Facility of the Haukeland University Hospital. We thank the entire Aquacosm-VIMS consortium for help and assistance in collecting, processing, and analyzing some of the core data used in this manuscript.

## Author Contribution

G.H., F.V. and A.V. conceived and wrote the paper. G.H. conducted all data analyses on SC-RNASeq. U.S. and C.K. collected and processed samples for SC-RNASeq. F.V performed ddPCR and smFISH experiments. All authors discussed the results and contributed to the final manuscript.

