## Supplementary Figures for "Daily turnover of active giant virus infection during algal blooms revealed by single-cell transcriptomics"

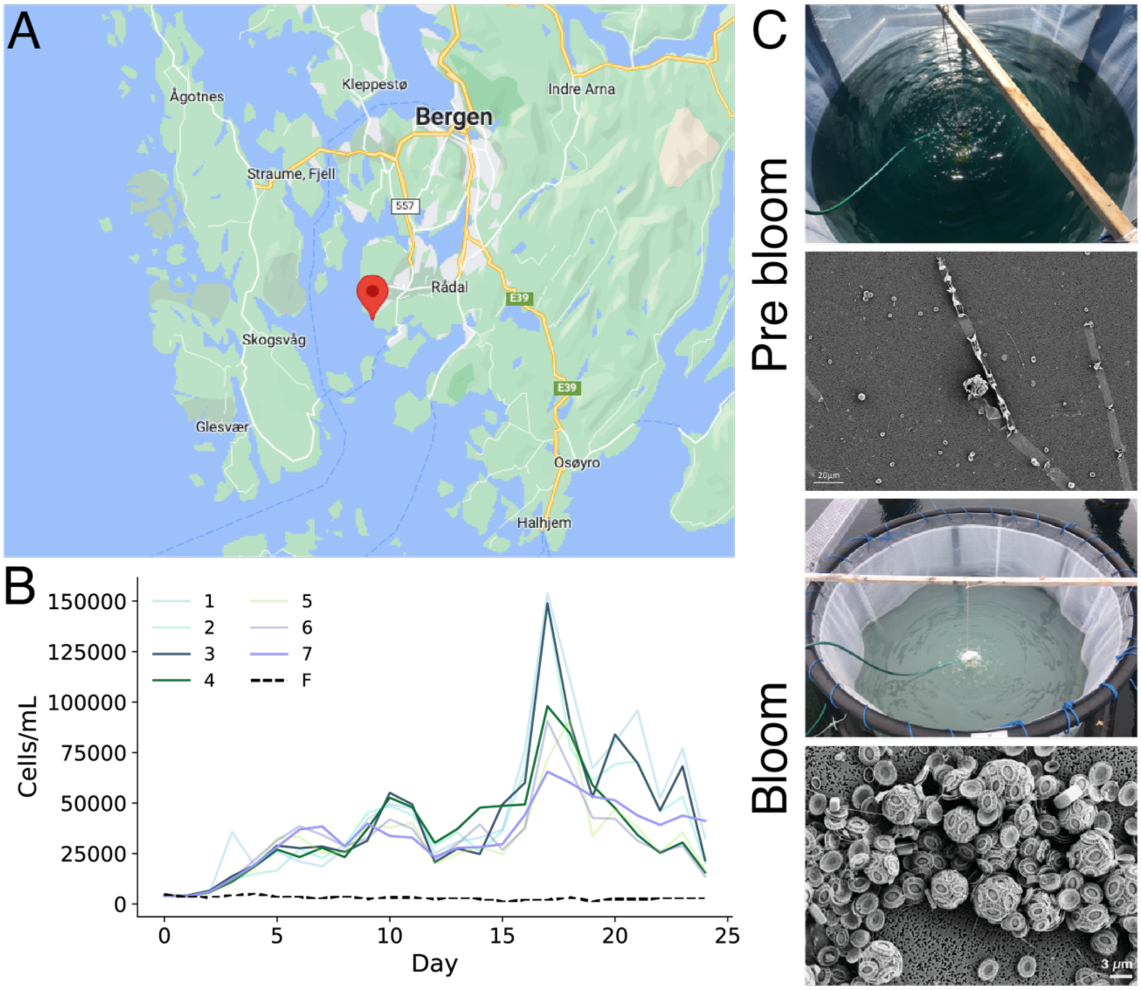


**Figure S1**. Mesocosm experiment in Norway. (A) Map of site in Norway (Google Maps). (B) Calcified *E. huxleyi* abundance measured by flow cytometry, based on high side scatter and high chlorophyll signals in the experiment bags (1-7) and the surrounding Fjord water. (C) Images of the water in the bags and under electron microscopy before and during the *E. huxleyi* bloom (images credit: Daniella Schatz and Michel J. Flores).

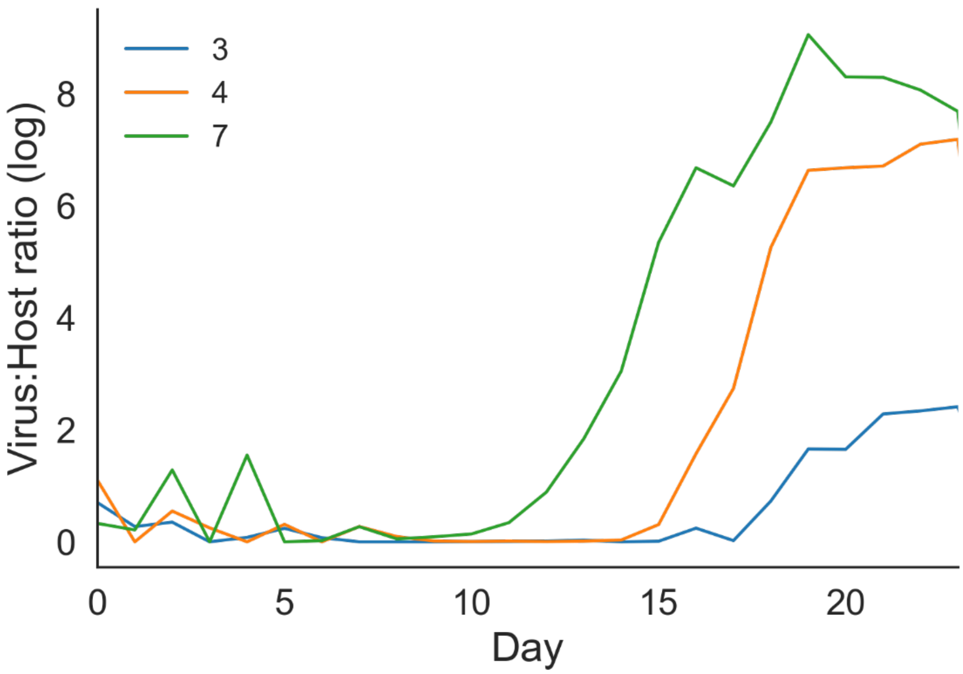


**Figure S2**. Virus to host daily ratios for bags 3, 4, and 7 throughout the mesocosm experiment. The *E. huxleyi* virus (EhV) abundance, estimated by qPCR using the major capsid protein gene *mcp* on biomass associated filters (2-20 μm).

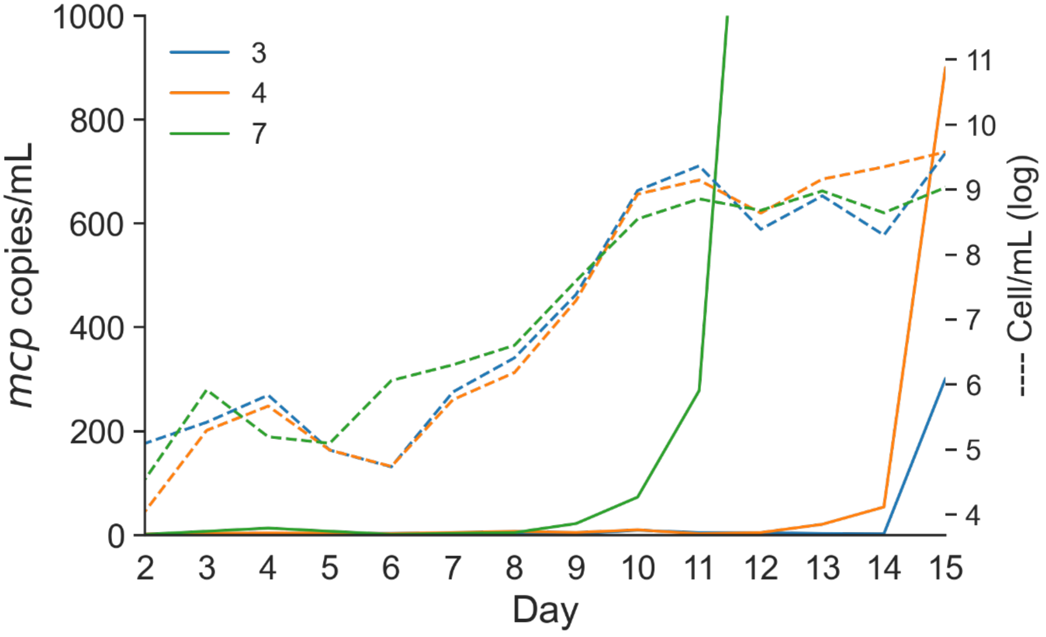


**Figure S3**. Early detection of viruses by ddPCR. Zoom in on the first days of the mesocosm experiment showing early detection of the major capsid protein gene *mcp* using digital droplet PCR to quantify intracellular viruses using 2-20 μm biomass filters (see methods). Virus abundance – solid lines. Host abundance – dashed lines.

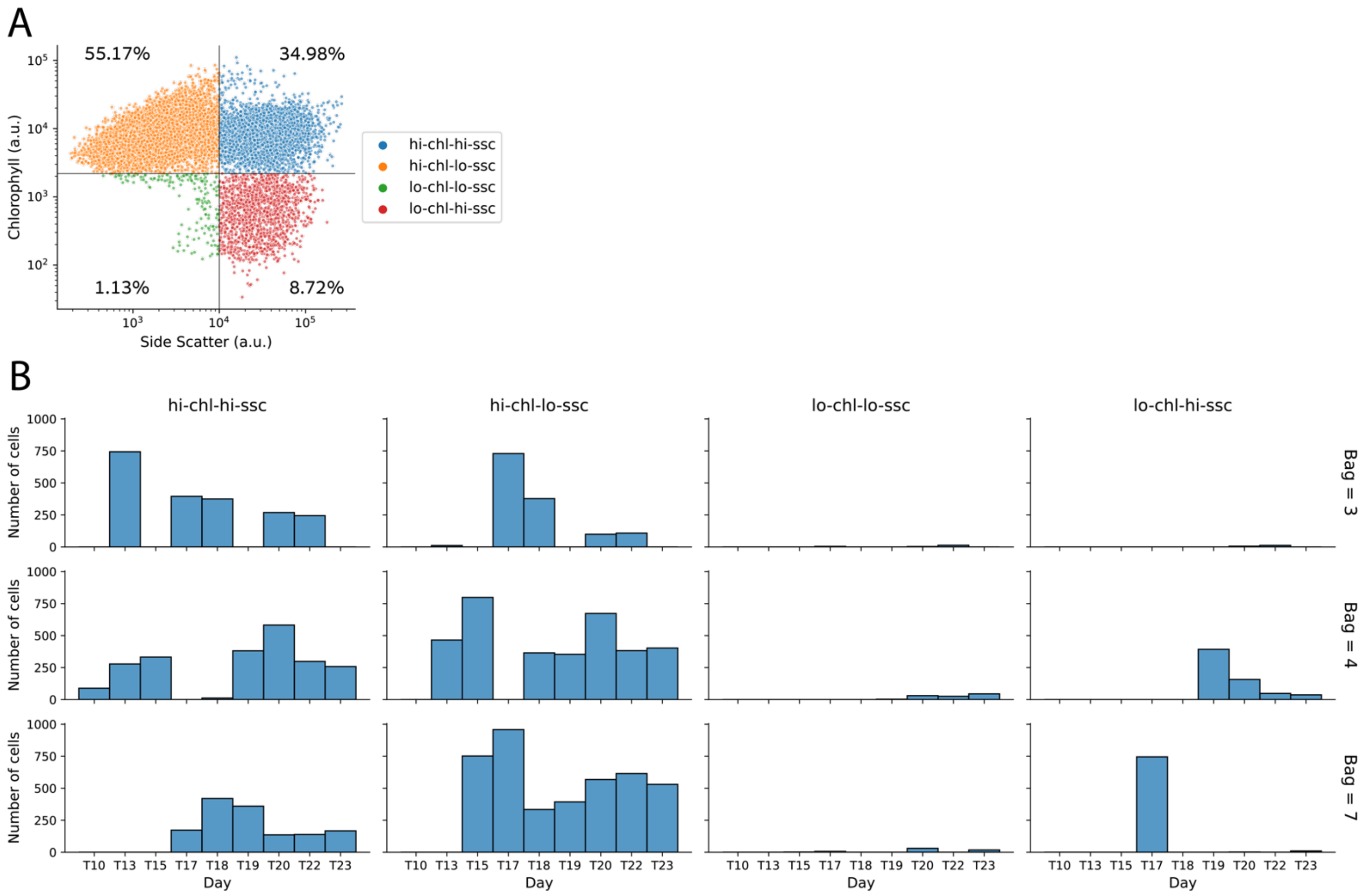


**Figure S4**. Single-cell FACS Index sorting for MARS-Seq scRNAseq. The sorting gates are defined based on the cells chlorophyll autofluorescence and side-scatter signal (A) All cells (N=12,769) distributed across the different sorting gates: Calcified *E. huxleyi* (high chlorophyll, high side-scatter), Naked *E. huxleyi* (high chlorophyll, low side-scatter), Calcified low chlorophyll *E. huxleyi* (low chlorophyll, high side-scatter), and non-target cells (low chlorophyll, low side-scatter). The percentage of cells from the total is annotated in each gate. (B) The number of sorted cells per bag-day-gate.

| Kinetic Class | Accession | Description | Details |
|---|---|---|---|
| Immediate-Early | AET98225.1 | hypothetical protein EPVG_00338 [Emiliania huxleyi virus 201] | https://www.ncbi.nlm.nih.gov/protein/AET98225.1/ |
| Immediate-Early | AET98234.1 | hypothetical protein EPVG_00347 [Emiliania huxleyi virus 201] | https://www.ncbi.nlm.nih.gov/protein/AET98234.1 |
| Immediate-Early | AET98238.1 | hypothetical protein EPVG_00351 [Emiliania huxleyi virus 201] | https://www.ncbi.nlm.nih.gov/protein/AET98238.1 |
| Immediate-Early | AET98219.1 | hypothetical protein EPVG_00332 [Emiliania huxleyi virus 201] | https://www.ncbi.nlm.nih.gov/protein/AET98219.1 |
| Late2 | AET97971.1 | major capsid protein [Emiliania huxleyi virus 201] | https://www.ncbi.nlm.nih.gov/protein/AET97971.1 |
| Late2 | AET98293.1 | hypothetical protein EPVG_00406 [Emiliania huxleyi virus 201] | https://www.ncbi.nlm.nih.gov/protein/AET98293.1 |
| Late2 | AET98065.1 | hypothetical protein EPVG_00178 [Emiliania huxleyi virus 201] | https://www.ncbi.nlm.nih.gov/protein/AET98065.1 |
| Late2 | AET97922.1 | hypothetical protein EPVG_00034 [Emiliania huxleyi virus 201] | https://www.ncbi.nlm.nih.gov/protein/AET97922.1 |
| Late2 | AET98248.1 | hypothetical protein EPVG_00361 [Emiliania huxleyi virus 201] | https://www.ncbi.nlm.nih.gov/protein/AET98248.1 |
| Late2 | AET98063.1 | hypothetical protein EPVG_00176 [Emiliania huxleyi virus 201] | https://www.ncbi.nlm.nih.gov/protein/AET98063.1 |
| Late2 | AET98323.1 | hypothetical protein EPVG_00436 [Emiliania huxleyi virus 201] | https://www.ncbi.nlm.nih.gov/protein/AET98323.1 |

**Table S1**. Infection state markers. Suggested list of highly expressed EhV marker genes for the infection states *Immediate-Early* (IE), and *Late2* (L2). See main text for details.
